# Local and dynamic regulation of neuronal glycolysis *in vivo*

**DOI:** 10.1101/2023.08.25.554774

**Authors:** Aaron D. Wolfe, John N Koberstein, Chadwick B Smith, Melissa L Stewart, Marc Hammarlund, Anthony Hyman, Philip JS Stork, Richard Goodman, Daniel A. Colón-Ramos

## Abstract

Energy metabolism supports neuronal function. While it is well established that changes in energy metabolism underpin brain plasticity and function, less is known about how individual neurons modulate their metabolic states to meet varying energy demands. This is because most approaches used to examine metabolism in living organisms lack the resolution to visualize energy metabolism within individual circuits, cells, or subcellular regions. Here we adapted a biosensor for glycolysis, HYlight, for use in *C. elegans* to image dynamic changes in glycolysis within individual neurons and *in vivo*. We determined that neurons perform glycolysis cell-autonomously, and modulate glycolytic states upon energy stress. By examining glycolysis in specific neurons, we documented a neuronal energy landscape comprising three general observations: 1) glycolytic states in neurons are diverse across individual cell types; 2) for a given condition, glycolytic states within individual neurons are reproducible across animals; and 3) for varying conditions of energy stress, glycolytic states are plastic and adapt to energy demands. Through genetic analyses, we uncovered roles for regulatory enzymes and mitochondrial localization in the cellular and subcellular dynamic regulation of glycolysis. Our study demonstrates the use of a single-cell glycolytic biosensor to examine how energy metabolism is distributed across cells and coupled to dynamic states of neuronal function, and uncovers new relationships between neuronal identities and metabolic landscapes *in vivo*.

**Significance statement:** While it is generally accepted that energy metabolism underpins neuronal function, how it is distributed and dynamically regulated in different tissues of the brain to meet varying energy demands is not well understood. Here we utilized a fluorescent biosensor, HYlight, to observe glycolytic metabolism at cellular and subcellular scales *in vivo*. By leveraging both the stereotyped identities of individual neurons in *C. elegans,* and genetic tools for manipulating glycolytic metabolism, we determined that neurons perform and dynamically regulate glycolysis to match changing cellular demands for energy. Our findings support a model whereby glycolytic states should be considered distinct and related to individual neuron identities *in vivo*, and introduce new questions about the interconnected nature of metabolism and neuronal function.

## Introduction

The brain utilizes large amounts of energy, much of which is derived from the metabolism of glucose via glycolysis and oxidative phosphorylation. In humans, the metabolic demands of discrete brain regions can be visualized using techniques such as fMRI or fPET, which reflect oxygen use and glucose uptake(1, 2, 8–11). Such measures show a strong correlation of increased metabolic demand with regions of neuronal activity(1–3, 9), indicating a link between brain activity and energy metabolism. While these studies suggest that energy metabolism responds to, supports, and flexibly adapts to brain activity states, it is not well understood how individual neurons modulate their metabolic states to meet the varying energy demands within the nervous system, as most approaches used to examine metabolism in living organisms lack the resolution necessary to visualize metabolic changes within individual circuits, single cells, or subcellular regions.

The distribution of glycolytic energy metabolism in the brain is particularly important. Glycolysis occurs in the brain under aerobic conditions, and increases in glycolysis have been associated with both plasticity and elevated neuronal activity(3–5). However, the extent to which neurons perform glycolysis remains controversial. One model posits that glycolysis is largely performed by support cells such as astrocytes(12–14), which cater to neuronal energy demands by shuttling lactate into adjacent neurons, where it can be converted to pyruvate and transported into mitochondria for oxidative phosphorylation. In this model, regulation of glycolysis and oxidative phosphorylation in the nervous system is distributed across cell types, with a net flux of lactate into neurons from support cells in response to neuronal stimulation. However, recent studies suggest neurons do increase glucose metabolism upon stimulation(5, 15–20). To reconcile these findings and better understand the role of glycolysis in energy metabolism across the nervous system, additional direct cell biological readouts of glycolytic activity across different cell types and in varying conditions are necessary.

Probing of energy metabolism with cellular and subcellular resolution benefits both from simpler models that enable single cell inspection *in vivo,* and tools that report on energy metabolic pathways at a single cell level. The *C. elegans* nervous system, with only 302 neurons and a well-understood connectome(21), satisfies the first requirement. Yet tools capable of directly monitoring glycolytic energy metabolism *in vivo* have been lacking. The genetically-encoded biosensor HYlight(7) was recently designed to monitor levels of fructose-1,6-biphosphate (FBP) as a proxy for levels of glycolytic activity. FBP is produced during glycolysis by the highly regulated enzyme phosphofructokinase (PFK), an enzyme that is allosterically inhibited by metabolites such as ATP and citrate (generated from the TCA cycle), activated by AMP and ADP, and modulated by other central metabolic regulators such as the proteins AMPK and PFKFB. Therefore, regulation of PFK—and thus regulation of FBP concentration—represents a convergence point of pathways responding to cellular energy states at the committed step of glycolysis. HYlight has been successfully used to monitor FBP levels in cultured pancreatic beta cells(7), a cell type that regulates glycolysis based on concentration of glucose in the medium. The sensor has not yet been used to monitor glycolytic states of tissues *in vivo*.

In this study we have adapted HYlight for use *in vivo* in *C. elegans*. We leveraged the stereotyped nervous system of this organism, the HYlight biosensor, and a microfluidic imaging device to characterize the glycolytic energy landscape in neurons at resting states and during energy stress. Using these approaches, we determined that neurons perform glycolysis cell autonomously, and that they dynamically modulate their glycolytic states. Taking advantage of the ability of HYlight to reveal metabolic differences across individual cells, we examined the state of glycolysis in specific neurons to uncover a complex neuronal and glycolytic energy landscape which revealed three main observations: 1) glycolytic states in neurons are diverse across individual cell types; 2) for a given condition, glycolytic states within individual neurons are reproducible across animals, and 3) for varying conditions of energy stress, glycolytic states are plastic and adapt to energy demands. Using a mutation that disrupts trafficking of mitochondria to synaptic regions, we showed that a lack of mitochondria near synapses resulted in increased localized glycolysis, thereby demonstrating subcellular differences in the regulation of glycolysis. Moreover, our genetic studies found that the *C. elegans* homologs of the glycolysis-regulating protein PFKFB are specifically required for adaptation of glycolysis in response to neuronal activity states, but not in response to transient hypoxia. Thus, our findings have uncovered multiple genetic regulators of neuronal glycolytic states that are responsible for adaptation to different sources of energy stress in *C. elegans*.

## Results

### Measuring neuronal glycolysis in vivo

To characterize the glycolytic profiles of neurons *in vivo*, we adapted the HYlight sensor for use in *C. elegans*. We achieved sufficient expression of the construct via codon optimization, introduction of introns derived from the *C. elegans* genome, and employment of cell specific promoters to drive stable expression of the sensor in desired tissues (Figures 1 and S1; Methods). As a control, we used a version of HYlight with reduced <u>a</u>ffinity for FBP that does not respond to changes in concentration of the FBP metabolite (HYlight-RA; Figure S1). Importantly, because HYlight is a ratiometric sensor, it provides a readout for FBP that is independent of protein expression. The ratiometric property of this sensor allowed us to compare FBP levels, as well as changes in FBP upon specific conditions and across different cells, tissues, or animals.

**Figure 1:**
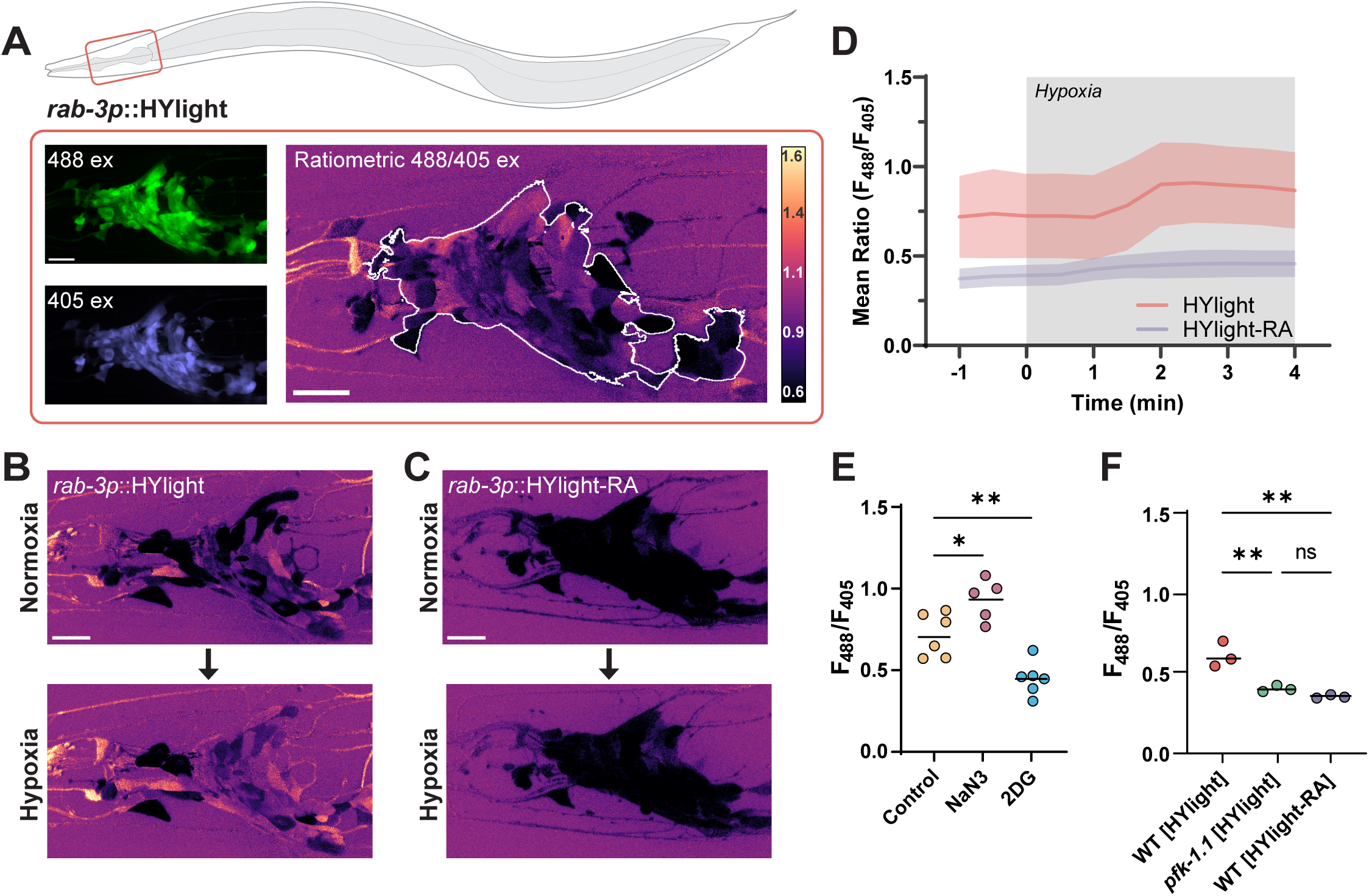
Measurements of glycolytic dynamics *in vivo* in *C. elegans* neurons. **(A)** Schematic of *C. elegans*, with imaged region highlighted by red box. The imaged area corresponds to the nerve ring (brain) of the nematode. HYlight and HYlight-RA have two excitation wavelengths that vary according to the level of FBP(7). Ratiometric images depict the ratio of 488 nm to 405 nm excitation of the biosensor. Scale bars, 10 μm. **(B)** Neuronal expression of HYlight (strain DCR8881) under normoxic (top) conditions, or after two minutes of hypoxic (bottom) conditions. **(C)** As in (B), but expressing HYlight-RA, a variant with reduced affinity for FBP used as a control. **(D)** Quantification of the mean ratio for the nerve ring of each worm shown in (B) and (C). Grey region of the graph denotes period in which the worms were exposed to hypoxic conditions, and shaded region around graphed lines denotes the standard deviation of the mean ratio at each time point. **(E)** Quantification of the HYlight ratio for worms treated with glucose (control), sodium azide (NaN3), or 2-deoxy-2-glucose (2DG) as described in Methods. Each dot corresponds to one worm; significance of * or ** denotes P values of 0.028 and 0.0051 respectively, as calculated by ANOVA with Tukey post-hoc test. Horizontal bar corresponds to the mean. **(F)** Quantification of HYlight or HYlight-RA ratios in wild-type and *pfk-1.1(ola458)* mutant animals; significance represents P-values of 0.0044 (WT vs *pfk-1.1*), 0.0015 (HYlight vs HYlight-RA), and 0.47 (*pfk-1.1* vs HYlight-RA), as calculated by ANOVA with Tukey post-hoc test.

Transient hypoxia reduces respiration by inhibiting mitochondrial Complex IV, decreasing cellular ATP levels, and is expected to increase the rate of glycolysis due to energy stress. Thus, manipulation of transient hypoxia in living animals allow us to directly control dynamics of glycolysis. To achieve this, we used a hybrid microfluidic-hydrogel device that enables precise regulation of oxygen concentration during high-resolution and long-term imaging of living *C. elegans* animals(22). We then generated transgenic animals that express neuronal HYlight or HYlight-RA by using the *rab-3* pan-neuronal promoter (Figure 1A). We observed that under normoxic conditions, HYlight displayed a broad range of ratios in different neurons, consistent with varying glycolytic states (and FBP levels) across cells (Figure 1B). In contrast, control HYlight-RA displayed low levels of signal in all cells (Figure 1C). Upon exposure to transient hypoxia, we observed an increase in the HYlight ratio within one to two minutes, which was not observed in animals expressing HYlight-RA (Figures 1B-D, Movie S1). These findings indicate that we can observe dynamic changes to glycolytic metabolism in neurons by using HYlight as a measurement of cellular FBP.

Next, to determine whether the observed changes in HYlight fluorescence corresponded to predicted manipulations of glycolytic flux, we used pharmacological and genetic approaches. We compared worms grown on plates containing either 2.5 mM 2-deoxy-2-glucose (2DG), an analogue of glucose that cannot be metabolized and blocks glycolysis, or 2.5 mM glucose as a control. When worms grown on glucose were exposed to sodium azide, an inhibitor of mitochondrial Complex IV, we observed a significantly increased mean HYlight ratio compared to the control worms, similar to the observed effects of transient hypoxia (Figure 1E; for experimental conditions used to minimize lethality, see Methods). By comparison, worms grown on 2DG, which inhibits glycolysis, displayed a significantly reduced mean HYlight ratio compared to control worms (Figure 1E).

FBP, the metabolite sensed by HYlight, is the product of the enzyme PFK. To further validate the observed ratio as a measurement of FBP, we generated, via CRISPR-Cas9, a knockout of *pfk-1.1*, the principal gene encoding PFK in somatic tissues of *C. elegans*(23). We observed that *pfk-1.1* knockout animals displayed reduced ratiometric values compared to wild type, and that these values were reminiscent of those seen for the HYlight-RA control (Figure 1F; for phenotypic description of the *pfk-1.1(ola458)* allele, see Methods). These findings indicate that the observed HYlight responses corresponded to expected increases or decreases in glycolytic metabolism and provides a sense of the dynamic range of the sensor for changes *in vivo*.

Given the ability of HYlight to detect a range of different glycolytic states, we next compared the level of glycolysis across different neuron classes (Figure 2A). For this, we took advantage of the invariant neurodevelopmental lineage of *C. elegans* to quantify glycolytic states in identified neurons across different animals. When imaged under similar conditions, the neurons ALN, DVB, and PLM displayed similar patterns of HYlight signals across different animals (Figure 2B and 2C). These findings suggest that there are cell-specific glycolytic states for individual neurons that are characteristic of neuronal identities.

**Figure 2:**
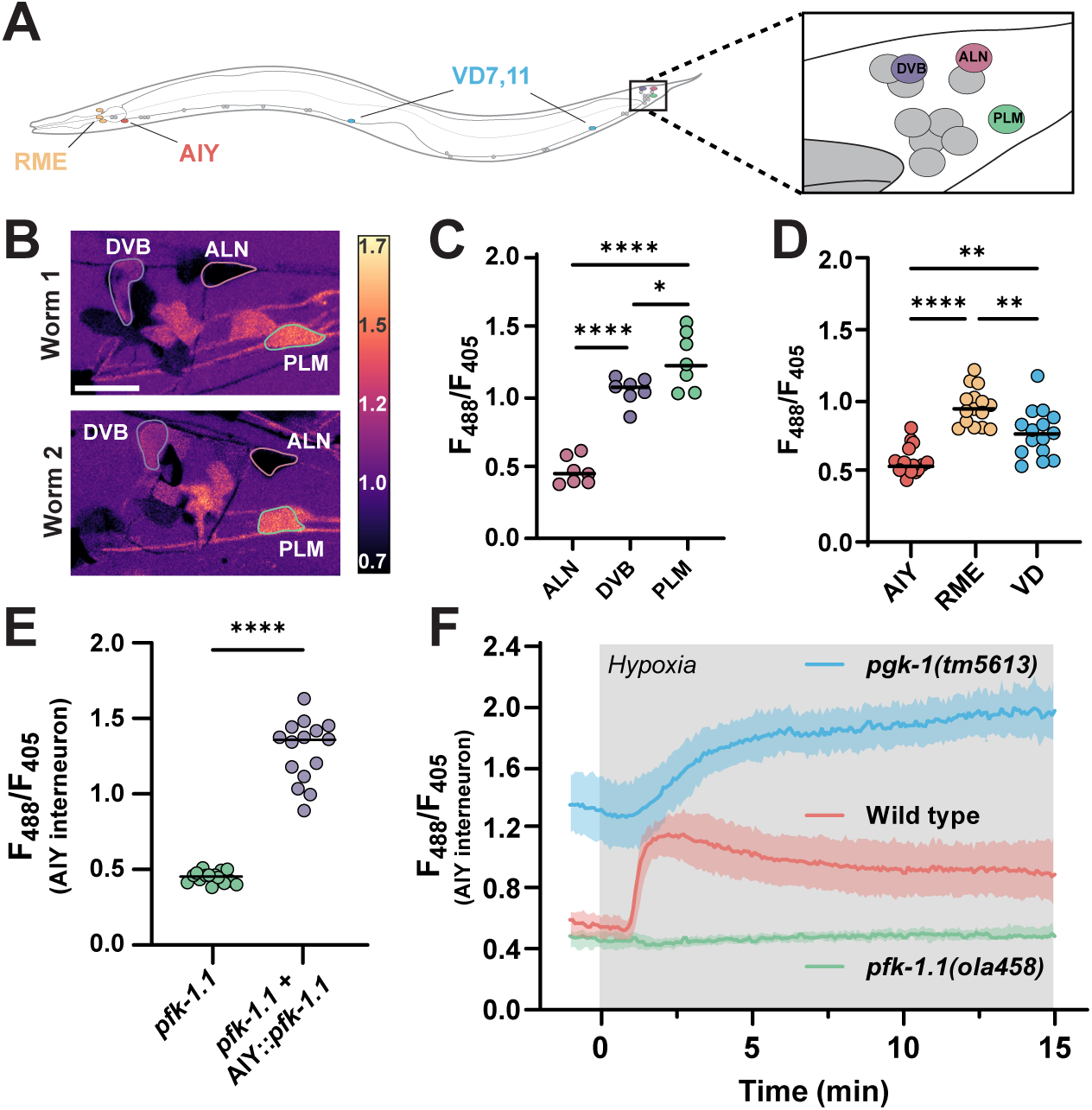
Glycolytic profiles map onto neuronal identities *in vivo*. **(A)** Schematic of *C. elegans* with positions of individual, identifiable neurons characterized in this study labeled, including tail ganglia (highlighted with box on the right) corresponding to images in (B). VD neurons were quantified using VD7 and VD11. **(B)** Pan-neuronally expressed HYlight, showing 488/405 nm ratio of excitation for neurons in the tail ganglia of two representative worms (Worm 1, top; Worm 2, bottom) under normoxic conditions, with identifiable neurons (DVB, ALN and PLM) labeled. Scale bar, 10 μm. **(C)** As (B), but quantified for seven individual worms. Significance values of * or **** represent P-values of 0.0299 or <0.0001, respectively, as calculated by ANOVA with Tukey post-hoc test. **(D)** Cell-specific promoters were used to express HYlight in either AIY (*ttx-3p*), or RME and VD neurons (*unc-47p*), with ratios quantified under normoxic conditions in 15 animals. Significance values are 0.0012 (AIY vs VD), <0.0001 (AIY vs RME), and 0.0018 (RME vs VD) via ANOVA with Tukey post-hoc test. **(E)** HYlight examined in AIY interneurons of *pfk-1.1(ola458)* mutant animals, or with cell-specific expression of wild-type PFK in AIY in this same mutant. Significance values indicate P-values of <0.0001 (****) as calculated by unpaired t-test for 15 animals. **(F)** HYlight responses in varying genetic backgrounds under transient hypoxia. There is an increase in glycolysis in wild-type worms within 2 minutes of hypoxia treatment (see also Movie S2). *pfk-1.1(ola458)* mutant animals start at lowered levels of HYlight signal (as compared to wild type) and fail to increase upon hypoxia. The loss-of-function mutation *pgk-1(tm5613)*, an enzyme downstream in the glycolytic pathway (Figure S2), starts at an elevated level of FBP and increases further upon transient hypoxia. Shading represents the standard deviation around mean values for 15 worms per treatment. Hypoxia treatment represented by grey box.

To examine this hypothesis further, we next labeled individual neurons by expressing HYlight in three distinct cell types—AIY, RME, and VD neurons—by using cell-specific promoters. We observed that these also had distinct and characteristic levels of glycolysis (Figure 2D), providing additional evidence that different neuron types display specific glycolytic profiles. We focused on the AIY interneurons to examine the effects of specific mutations in glycolytic enzymes. The *pfk-1.1(ola458)* deletion significantly reduced the ratio compared to wild type (Figure S2), as observed with pan-neuronal HYlight expression (Figure 1E). Cell-specifically rescuing this mutation by expressing the wild-type PFK cDNA in AIY (*ttx-3p::pfk-1.1*) prevented this effect and increased the HYlight ratio (Figure 2E).

To follow dynamic changes of glycolytic flux in single cells during transient hypoxia, we examined HYlight responses in AIY neurons and in the context of mutations that disrupt either the upstream or downstream glycolytic pathways (Figures 2F and S2). In wild-type worms we observed increased HYlight responses that occurred within 1-2 minutes of initiating the hypoxia treatment (Figure 2F, Movie S2). This increase failed to occur in the *pfk-1.1* mutant. Conversely, a mutation in the gene *pgk-1,* which is downstream of *pfk-1.1* in the glycolytic pathway, displayed elevated ratios of HYlight responses as compared to wild type, consistent with the expected decreased utilization of FBP by downstream steps of the pathway. These observations indicate that HYlight measurements directly represent expected dynamics resulting from glycolytic flux through PFK in neurons, and support that neurons can perform glycolysis *in vivo*.

### Pharmacologically altering neuronal activity affects the level of glycolysis

To test whether pharmacological agents capable of activating or silencing neurons affect the level of glycolysis, we examined the effects of levamisole and muscimol on HYlight measurements. Levamisole is an agonist of nicotinic acetylcholine receptors and activates neurons expressing these receptors. Muscimol is an agonist of ionotropic GABAa receptors and silences neurons with this receptor type. We exposed worms expressing pan-neuronal HYlight to either levamisole or muscimol, and quantified HYlight responses. Animals exposed to levamisole displayed elevated ratios of HYlight values as compared to animals exposed to muscimol (Figure 3A). We next compared the level of glycolysis under these two pharmacological treatments, but specifically in VD neurons. VD neurons express both nicotinic acetylcholine receptors and ionotropic GABAa receptors and should be directly affected by these reagents(23–26). When worms were treated with levamisole, we observed a significant increase in the level of glycolysis in VD neurons as compared to a buffer control, reflecting an increase of glycolytic state upon persistent neuronal depolarization (Figure 3B). In contrast, when worms were exposed to muscimol, we observed a significant decrease in the HYlight ratio, reflecting the reduction in energy demand that occurs upon neuronal silencing (Figure 3B).

**Figure 3:**
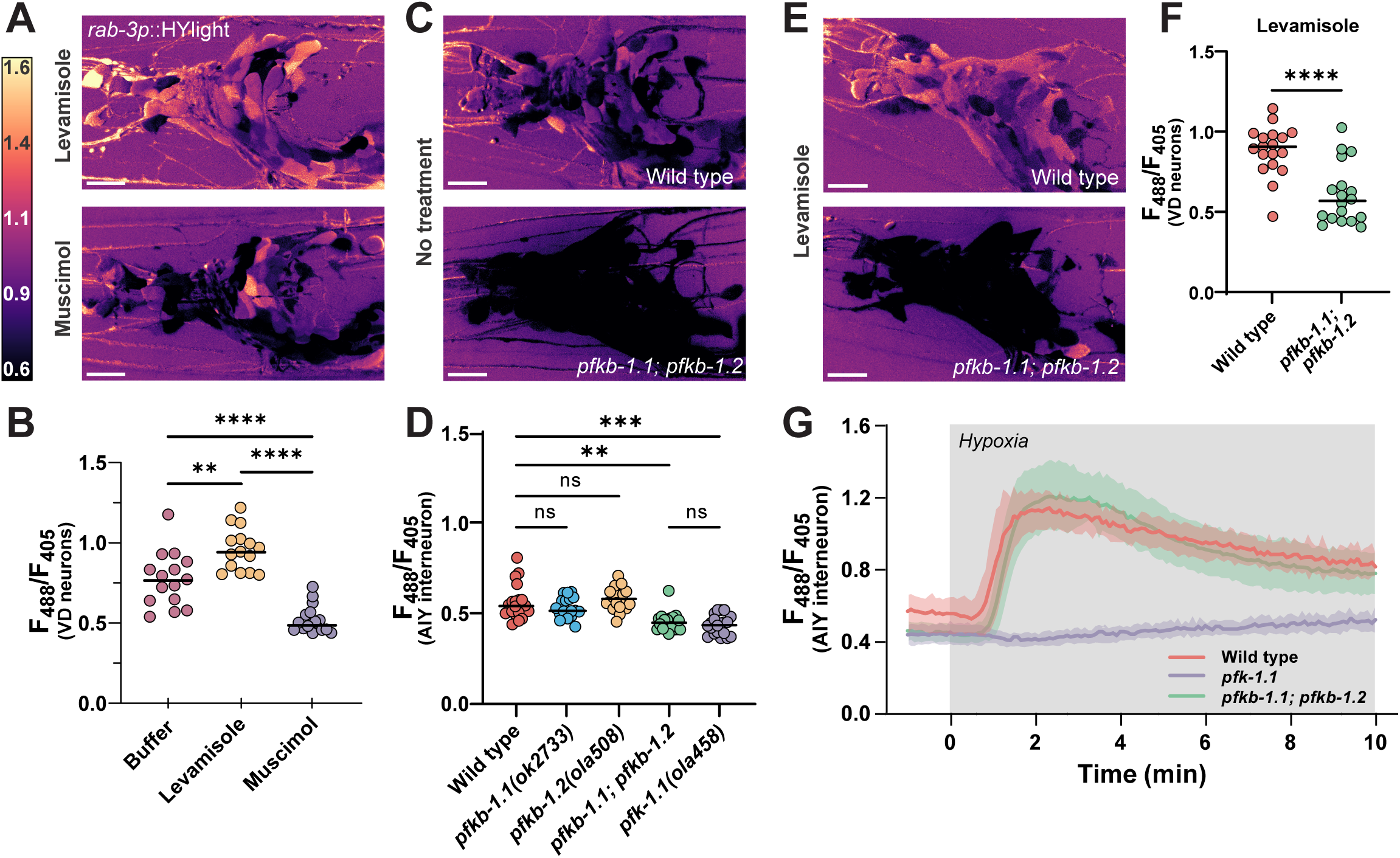
Neurons depend on PFKFB enzymes to modulate rates of glycolysis. **(A)** HYlight ratio of 488/405 nm excitation in the nerve ring of worms mounted in 10 mM levamisole (top) or 50 mM muscimol (bottom). Scale bar, 10 μm. **(B)** Quantification of HYlight ratios in the VD neurons when worms are mounted in either M9 buffer, 10 mM levamisole, or 50 mM muscimol. Significance represents P-values of 0.0013 (**) or <0.0001 (****), as calculated by ANOVA with Tukey post-hoc test, for 15 animals. **(C)** HYlight ratios in the nerve ring of wild type (top) or *pfkb-1.1(ok2733); pfkb-1.2(ola508)* double mutants (bottom) mounted in M9 buffer. Single mutant images and quantification in Figure S3. **(D)** Quantification of HYlight responses for indicated genotypes in the AIY interneuron. Significance represents P-values of 0.512 (WT vs. *pfkb-1.1*), >0.999 (WT vs. *pfkb-1.*2), 0.927 (WT vs. *pfkb-1.1; pfkb-1.2*), or <0.0001 (**), as calculated with ANOVA/Tukey post-hoc test across 18 animals. **(E)** HYlight ratio in the nerve ring of wild-type and *pfkb-1.1(ok2733); pfkb-1.2(ola508)* double mutant animals exposed to 10 mM levamisole. **(F)** HYlight ratios quantified for indicated genotypes in VD motor neurons in the presence of 10 mM levamisole. P-value of <0.0001 (****) is shown as calculated by unpaired t-test across 18 animals. **(G)** HYlight responses in the AIY interneuron for the indicated genotypes under transient hypoxia. The *pfk-1.1(ola458)* mutant animals do not increase upon hypoxia, whereas *pfkb-1.1(ok2733); pfkb-1.2(ola508)* double mutant animals, which start at a similar mean ratio as shown in (D), display responses to transient hypoxia that are indistinguishable from wild type. Shading represents the standard deviation around mean values for 18 worms per treatment. Hypoxia treatment represented by grey box.

### PFKFB enzymes are necessary for neuronal glycolysis

In mammalian neurons, glycolytic flux is controlled to a large degree by fructose-2,6-bisphosphate (F2,6BP), which blocks the suppressive effect of ATP on PFK. F2,6BP is produced by PFKFB, which has two orthologs in *C. elegans*, *pfkb-1.1* and *pfkb-1.2*, both widely expressed in neurons(23). As no allele for *pfkb-1.2* was yet present in literature, we used CRISPR-Cas9 to generate a full deletion of the gene (see Methods). A comparison of pan-neuronal HYlight in single knockouts of either PFKFB gene, *pfkb-1.1(ok2733)* and *pfkb-1.2(ola508)*, showed no change in the mean level of glycolysis; however, a double knockout reduced glycolysis significantly (Figure 3C and S3). This same result was observed in AIY, which expresses both genes(23)—there was no significant difference between the level of glycolysis in either single mutant compared to wild type, whereas the double mutant was reduced to the same level as a mutant of *pfk-1.1* (Figure 3D). This suggests that PFKFB enzymes are necessary for glycolysis to occur in neurons, and that the two orthologues of PFKFB in *C. elegans* (*pfkb-1.1* and *pfkb-1.2*) are functionally redundant in *C. elegans*.

We next asked whether the elevated HYlight signal observed after treatment with levamisole also occurred in the absence of PFKFB. We examined wild-type or *pfkb-1.1; pfkb-1.2* double mutant worms expressing pan-neuronal HYlight and observed that the double-mutant animals had a broadly reduced HYlight ratio despite the levamisole treatment (Figure 3E). To quantify this, we repeated this experiment in worms expressing HYlight in VD neurons, for which we similarly observed that the HYlight ratio in the double *pfkb-1.1; pfkb-1.2* mutant background was significantly reduced compared to wild-type worms (Figure 3F). This indicates that the increase in glycolysis induced by levamisole treatment cannot occur in the absence of PFKFB, and likely requires F2,6BP for this increase to occur. Interestingly, when worms expressing HYlight in the *pfkb-1.1; pfkb-1.2* mutant background were exposed to hypoxia, the rise in FBP level still occurred (Figure 3G). Our genetic studies suggest that increases in glycolysis in neurons due to different conditions (hypoxia vs neuronal activity) are regulated by distinct mechanisms with varying dependencies on the PFKFB enzymes and regulation by F2,6BP.

### Subcellular changes in neuronal glycolysis relative to mitochondria

To better understand the interplay between respiration and glycolysis, we next tested HYlight responses in neurons of animals with defective mitochondrial function or localization. In a hypomorph mutant of mitochondrial complex III, *isp-1(qm150)*, FBP levels were elevated compared to wild-type worms under normoxic conditions and these levels did not increase further upon hypoxia treatment (Figure 4A). Thus, reduction of oxidative phosphorylation and inhibition of mitochondrial respiration consistently shifted metabolism towards elevated glycolysis, whether it was achieved via transient hypoxia, pharmacological inhibition, or genetics.

**Figure 4:**
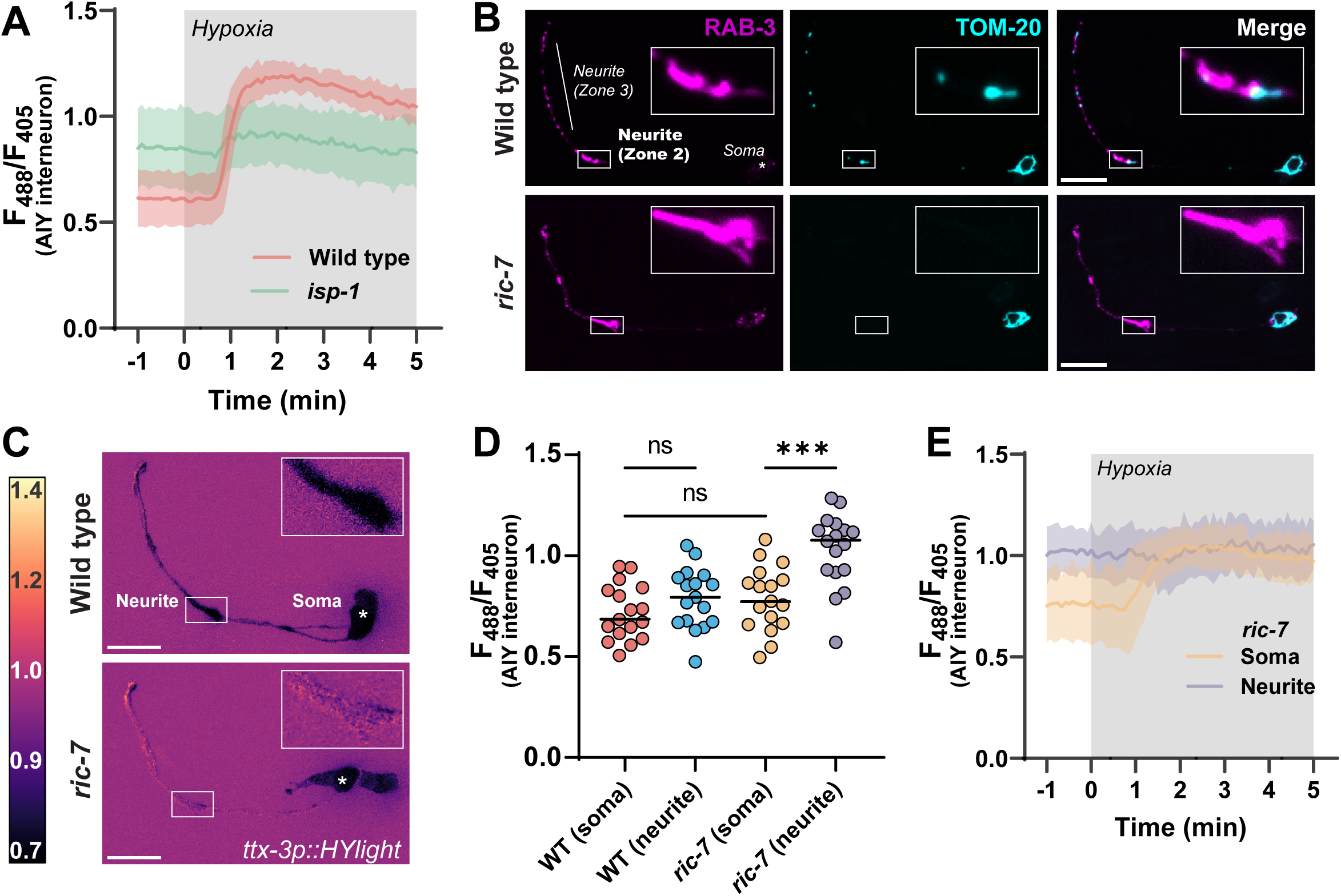
Local regulation of glycolysis in neurons upon disruptions of mitochondria localization. **(A)** HYlight responses in a mutant of mitochondrial complex III, *isp-1(qm150),* and wild-type animals, upon transient hypoxia in the AIY interneuron. Glycolysis is elevated in *isp-1* mutant animals prior to hypoxia treatment, and show reduced glycolytic response to hypoxia compared to wild-type worms. The shaded region per each graphed line represents standard deviation of the mean of 11 animals; hypoxia treatment is highlighted in grey. **(B)** Localization of synaptic vesicles (first column) and mitochondria (second column) in the AIY interneurons of wild type (first row) and *ric-7(n2657)* mutants (second row). AIY has distinct synaptic regions in the neurite(21, 41), labeled as Zone 2 (inset) and Zone 3, near which mitochondria localize in wild-type animals. In a mutant of the kinesin adapter protein *ric-7*, mitochondria are incapable of localizing to synapses and are only found within the cell body, consistent with prior work(30, 31). Scale bar, 10 μm. **(C)** HYlight responses in wild-type (top) and *ric-7(n2657)* mutant animals (bottom) in the AIY interneurons under normoxic conditions. In *ric-7* mutants, Zone 2 of the neurite (inset) has a distinctly higher ratio compared to the cell soma, whereas these two regions are similar in wild-type worms. **(D)** Quantification of the HYlight ratio in different subcellular regions of AIY interneurons of wild-type and *ric-7(n2657)* mutant animals. P-values shown are 0.5568 (WT soma vs. neurite), 0.5835 (WT soma vs. *ric-7* soma), and 0.0002 (*ric-7* soma vs. neurite) as calculated by ANOVA with Tukey post-hoc test for 17 animals. **(E)** Subcellular responses to transient hypoxia in AIY interneurons in *ric-7(n2657)* mutant animals. The synaptic regions of AIY have elevated glycolysis prior to hypoxic treatment and do not respond upon treatment; the cell soma responds to hypoxia. Shading on each graphed line represents standard deviation of the mean for 13 animals; hypoxia treatment in grey box.

There is considerable evidence that different regions within a neuron have distinct metabolic capabilities depending on their differing and often dynamic energy requirements(27–29). Furthermore, the observed link between elevated glycolysis and mitochondrial dysfunction suggests that there may be subcellular differences in glycolysis depending on whether mitochondria are locally available. To address this idea, we utilized a mutant of a mitochondrial kinesin adapter protein, *ric-7(n2657)*, a protein that is necessary for transport of mitochondria to neurites(30, 31). In this mutant, mitochondria are trapped within the cell soma and excluded from neurite regions (Figure 4B). We observed that in this background HYlight displayed locally elevated glycolytic levels specifically within neurites (Figure 4C), while no significant differences were observed at the cell body of *ric-7* worms as compared to the cell body of wild-type neurons (Figure 4D). Moreover, hypoxia treatment of *ric-7* mutant worms showed that, while HYlight ratios within the cell soma increased upon hypoxia treatment, no further changes were observed in the elevated neurite regions (Figure 4E). Thus, disrupting mitochondria trafficking to nerve terminals caused a localized shift to increased glycolysis within the neurite. Our findings demonstrate a capacity for subcellular regulation of glycolytic flux within different compartments of the neuron.

## Discussion

While it is generally understood that glucose is a critical fuel for neuronal function, the regulation of neuronal glycolysis remains controversial(14, 19). At one extreme, studies have argued that neurons have a limited capacity to perform glycolysis, and that astrocyte-derived lactate is the principal fuel for neuronal ATP. At the other, it is argued that neurons principally import and metabolize glucose through glycolysis, particularly in response to stimulation. Sorting out which of these models is correct requires tools capable of monitoring glycolysis *in vivo* at the single cell level. Here we establish such a system in *C. elegans* by using the FBP biosensor, HYlight. HYlight has enabled *in vivo* examination of the dynamics of glycolysis within individual neurons and upon varying physiological stimuli. Using this system, we have observed that 1) neurons perform glycolysis *in vivo*; 2) varying glycolytic states map to specific neuronal identities, and 3) neurons are capable of dynamically adapting their glycolytic states, even in subcellular regions, to meet energy demands.

HYlight is a powerful tool to examine glycolytic dynamics in single cells and across tissues. As a ratiometric sensor, it provides a measurement for the metabolite FBP that is largely independent of sensor expression, allowing for comparisons across different cells, tissues, or individual animals. By making use of a nonbinding control variant, as well as key mutants of the glycolytic pathway, we were able to validate our HYlight measurements at the single cell level and append meaning to different ratiometric values, despite the challenges of working *in vivo.* Through this we were able to determine that different identifiable cells favor distinct levels of glycolysis under a particular condition, and that these characteristic levels of glycolysis are consistent when compared across individual animals. Importantly, this suggests that the metabolic profile of a neuron is defined in part by its identity, which may be due to different protein expression profiles, cellular environments, or functional roles within circuits.

Functional imaging studies have demonstrated that distinct metabolic states of brain regions are associated with both elevated neuronal activity and plasticity(1, 2, 11). Moreover, recent studies from *Drosophila* using pyruvate, ATP, and calcium sensors proposed that cells are capable of predictive energy allocation, in which metabolism is regulated to anticipate and meet energy demands based on neuronal activity states and history(6). It is therefore possible that the observed neuron-specific signatures of glycolytic metabolism reported in this study reflect typical activity or firing states for individual neurons. Precisely how glycolysis aligns with these states would need to be examined in future studies. Nonetheless, our findings uncover a reproducible glycolytic landscape across neuronal cell types *in vivo*, and opens up new opportunities to examine how these landscapes are regulated to meet neuronal function.

Neurons are capable of local regulation of glycolysis *in vivo*. Our findings demonstrate that while specific neurons display different levels of glycolysis under particular conditions, they are also capable of dynamically modulating glycolytic flux upon energy stress. This is consistent with prior work that observed changes in glucose or NADH to indirectly measure increases in neuronal glycolysis in an activity-dependent manner(4, 16, 32). Upon stimulation, increased levels of glucose are taken up by neurons preferentially within synaptic regions, and cellular NADH/NAD+ ratio increases in a manner indicative of elevated glycolysis(16). In our study we have extended these observations by directly visualizing a key rate-limiting metabolite of the glycolytic pathway *in vivo*, which provides an effective proxy for measuring glycolysis. By using microfluidic devices to modulate the environmental conditions of a worm, as well as genetic and pharmacological manipulations, we have shown that neurons can perform, and plastically regulate, glycolytic states to meet changing energy demands in a rapid and dynamic manner.

A double mutant of *pfkb-1.1* and *pfkb-1.2* dramatically reduced glycolysis in neurons. These two genes encode the only known homologues in *C. elegans* for PFKFB, a key regulator of glycolysis in vertebrates. PFKFB has dual kinase/phosphatase activity and modulates levels of the metabolite F2,6BP, which promotes glycolysis by blocking the inhibitory effects of ATP on PFK. It has been proposed that neuronal glycolysis in the mammalian brain is suppressed compared to astrocytic glycolysis and via the persistent degradation of the brain-enriched PFKFB3 paralog(33, 34). In this study, we demonstrate that the two *C. elegans* PFKFB orthologs are functionally redundant; but necessary to regulate glycolysis in neurons, particularly under conditions associated with increased neuronal activity such as after treatment with levamisole. Vertebrate cells similarly express other PFKFB paralogues(34), and we speculate that these could be important for regulating neuronal glycolysis. Interestingly, we did not observe a phenotype for *pfkb-1.1*; *pfkb-1.2* double mutants when glycolysis increased during transient hypoxia, indicating a genetic separation of the mechanisms that regulate different increases of glycolysis in neurons. We hypothesize this is due to the role of F2,6BP in counteracting ATP inhibition of PFK, and reflects a difference in the strength of energy deficits. Importantly, our findings suggest that neurons rely on different genetic pathways to address different energy stress conditions.

Neuronal energy metabolism, consisting primarily of glycolysis and oxidative phosphorylation, is organized at both cellular and subcellular scales. The idea that glycolysis is compartmentalized within neurons has considerable support. For example, GLUT4 is locally transported to presynapses upon stimulation and is necessary for local glucose import in support of exocytosis(15). Our lab documented the subcellular localization of glycolytic proteins to presynaptic sites, with specific enzymes such as PFK becoming enriched in synaptic condensates upon energy stress(22, 35). Subcellular localization of glycolytic enzymes has also been observed at the leading edge of anchor cells during invasion of the uterine basement membrane of *C. elegans*(36). While these observations are consistent with the regulation of local glycolytic metabolism at synapses, they do not prove that glycolysis is occurring in this region; first, because glucose uptake does not necessarily lead to glycolysis; and second, because the biochemical function of glycolytic metabolons *in vivo* has not been demonstrated. To examine local regulation of metabolism, we examined effects of a mutation that disrupts trafficking of mitochondria to synaptic regions. Failed transport of mitochondria resulted in increased FBP production in neurites, consistent with changes to localized glycolysis. This finding provides additional evidence for local regulation of energy metabolism in neurons. Our studies also set the stage for future efforts to uncover dynamic mechanisms that regulate local energy metabolism in neurons and near synaptic regions.

Energy metabolism underpins neuronal function, but the mechanisms that link energy metabolic pathways with neuronal function at the cellular level remain poorly understood. Our study has demonstrated the use of a single cell glycolytic biosensor to examine the cellular and subcellular regulation of metabolism, and has presented new relationships between neuronal identities and glycolytic energy landscapes *in vivo*.

## Methods

### C. elegans strains and genetics

All worm strains were raised on nematode growth media at 20°C using the *Escherichia coli* strain OP50 as the sole food source on NGM/agar plates(37). Analyses were performed using hermaphrodites in the L4 larval growth stage. The *C. elegans* Bristol strain N2 was used as wild-type controls. Strains used within this study can be found in Table S1. The PFK-1.1 knock-out allele generated for this study, *pfk-1.1(ola458)*, displayed embryonic lethality and slower developmental dynamics. Animals that developed to the L4 stage were used for analyses with HYLight. The PFKB-1.2 knock-out allele generated here, *pfkb-1.2(ola508)*, appeared superficially wild type, but the double mutant of *pfkb-1.1(ok2733) I; pfkb-1.2(ola508) IV* was phenotypically similar to the *pfk-1.1(ola458)* allele.

### Molecular Biology

Standard molecular biology methods were used to generate all constructs. Plasmids were derived from vector backbones originating from either pSM or pDEST variants and amplified from transformed *E. coli* strains containing a single plasmid. The HYlight biosensor was ordered as a GeneArt String dsDNA fragment from ThermoFisher; the construct was codon-optimized and a single intron was added at nucleotide 150 to improve expression(38). The HYlight-RA construct was generated by adding the T67E mutation known to disrupt F1,6BP binding(7). The plasmids expressing HYlight and HYlight-RA in all neurons (pDACR3882 and pDACR3883, respectively) are available on Addgene. Other plasmids and sequences used within this study are available upon request.

### Transgenics

Transgenic strains were made by standard germline injection techniques; injection mixes of plasmids were normalized to 100 ng/uL of total DNA concentration using an empty *E. coli* expression plasmid as filler. Stably transmitting strains that transmitted to the F2 generation were selected. The strain DCR8981 (*olaEx5367*) was integrated mapped and outcrossed to generate DCR9089 (*olaIs138 IV*), (strains EG8040 and XE1763 were used to map the integration). CRISPR-Cas9 was used to generate genetic knockouts for *pfk-1.1(ola458)* and *pfkb-1.2(ola508)* based on protocols previously described(39). For both alleles, the CRISPOR tool(40) was used to select gRNA sites adjacent to the 5’ and 3’ sites for each gene and ordered from Horizon Discovery as modified crRNA oligos. CRISPR editing was done by injecting Cas9 protein (Horizon) at 3 μM, that was preincubated with trRNA (40 μM, Horizon) at 37C for 10 minutes with both crRNAs (30 μM each) and a ∼150 base ssDNA repair template (0.5 μM, Keck Oligo Core with gel extraction purification) that removes the respective gene, with 30 bases of overlap to genomic sequence at either end; KCl and Hepes pH 7.3 was added to 25 and 7.5 mM, respectively. The injection mix also contained the appropriate crRNA (12 μM) and template for recreating the *dpy-10(cn64)* allele (0.25 μM) as a reporter. Worms from the F1 generation with the *roller* phenotype were singled and sequenced for the novel allele and the resulting homozygous progeny were outcrossed three times prior to use to remove the *dpy-10* allele as well as any background mutations.

### Microfluidics and mounting

Worms were imaged using a PDMS microfluidics device that allowed control of atmospheric conditions experienced while imaging as described previously(22). A pad of 10% agarose dissolved in water of approximately 0.06 mm thickness was applied to the top of the PDMS device. For imaging HYlight, a 2.5 μL drop of M9 buffer, or either 10 mM levamisole or 50 mM muscimol dissolved in M9 (as listed in figure legends), was placed onto the pad. 1-3 worms (pan-neuronal imaging) or 10-20 worms (single neuron imaging) at the L4 larval stage were then moved into the drop. A whisker pick was used to wick the buffer outwards and then a size 1.5 cover slip was placed on top. Air was flowed at approximately 7-8 deciliters/second through the device to prevent hypoxic conditions from occurring beneath the coverslip(35).

### Experiments with metabolic inhibitors

For treatment with metabolic inhibitors, adult worms expressing pan-neuronal HYlight (DCR8881) were placed on 6 cm plates containing either 2.5 mM 2-deoxy-2-glucose (2DG) or 2.5 mM glucose dissolved within the NGM base. F1 progeny with the array were selected at the developmental larva stage 4 for imaging. 2DG treatment decreased the number of progeny, and worms from multiple plates were combined for experiments. For microscopy, worms from the glucose plates were mounted as described above in M9 buffer, either with or without the addition of 5 mM sodium azide, and incubated for 15 minutes prior to imaging. While prolonged exposure to sodium azide results in lethality, transient exposure in the order of 15 minutes and as used in this study have been previously used in the field to anesthetize animals.

### Confocal microscopy

Microscopy was done using a Nikon Ti2 + CSU-W1 spinning disk confocal microscope. After slide mounting, a 5-minute equilibration period was observed to allow the slide to thermally equilibrate with the slightly elevated ambient temperature on the microscope. Images were captured with a Hamamatsu Orca-Fusion BT CMOS camera at 16-bit pixel depth. Excitation of samples was done using 50 mW lasers at the 405, 488, or 561 nm wavelengths as necessary. Ratiometric imaging of HYlight was performed by switching between the 488 and 405 nm lasers without changing the 525 nm emission filter; laser power was set to 8%/2% (for 10x imaging) or 8%/4% (for 60x imaging) respectively, with the same exposure time (10-100 ms) for both channels. Exposure time was modified to adjust for brightness of different lines as necessary, but was always kept equal in both channels. Imaging settings were calibrated for different objectives or lines with the HYlight-RA construct to give an average ratio of 488/405 excitation of ∼0.40. For hypoxia experiments, flow was changed manually between tanks containing either compressed breathing air or pure N2 gas (beginning at the times marked on charts).

### Data analysis

All images were processed using Fiji/ImageJ. For visualization, ratiometric images were prepared using the Image Calculator>Divide command to divide the 488 nm excitation channel by the 405 nm channel. LUT was set to mpl_magma and scaled to the ratio ranges shown per figure. Quantification of 60x panneuronal HYlight images was done with a script (available online at the Colón-Ramos lab Github: https://github.com/colonramoslab/Wolfe-2023_scripts) that, per Z slice, applies background subtraction with a 50-pixel rolling ball algorithm, applies a 1-sigma gaussian filter to both channels, creates an additive image of the two channels together, thresholds this image using the “Huang” algorithm, and creates an ROI of this threshold; the resulting ROI is then used to capture the pixel values of the corresponding ratiometric image from the same slice. These pixel values were exported and quantified in the resulting charts. For 10x captures, images were background subtracted as above, and a square ROI was drawn around the resulting region of interest (generally the cell soma, or the neurite as described in figures) per time frame. The mean pixel value of this ROI was then used as the representative value for this cell, and multiple cells from the same image were aggregated for data analysis.

## Acknowledgements

We thank current and former members of the D.C.-R. lab for advice, suggestions, and guidance, in particular Milind Singh, Snusha Ravikumar and Ian Gonzalez. We also thank Gill Pollmeier and Henrick Bringmann (Technische Universität Dresden, Germany), and Hadas Dabas (Hammarlund lab, Yale University) for helpful discussions on the reagents and metabolic pathways. We thank the Caenorhabditis Genetics Center (P40 OD010440) and the Mitani lab (Tokyo Women’s Medical University School of Medicine) for strains. We thank the Keck Oligo Synthesis Resource at Yale for their assistance with synthesis of short single-stranded DNA oligos. This work was funded by the grants DP1 NS111778 (National Institute of Neurological Disorders and Stroke) and RGP0023/2019 (Human Frontier Science Program) awarded to D.C.-R and A.H.

