## Supplemental Info for "Local and dynamic regulation of neuronal glycolysis *in vivo*"

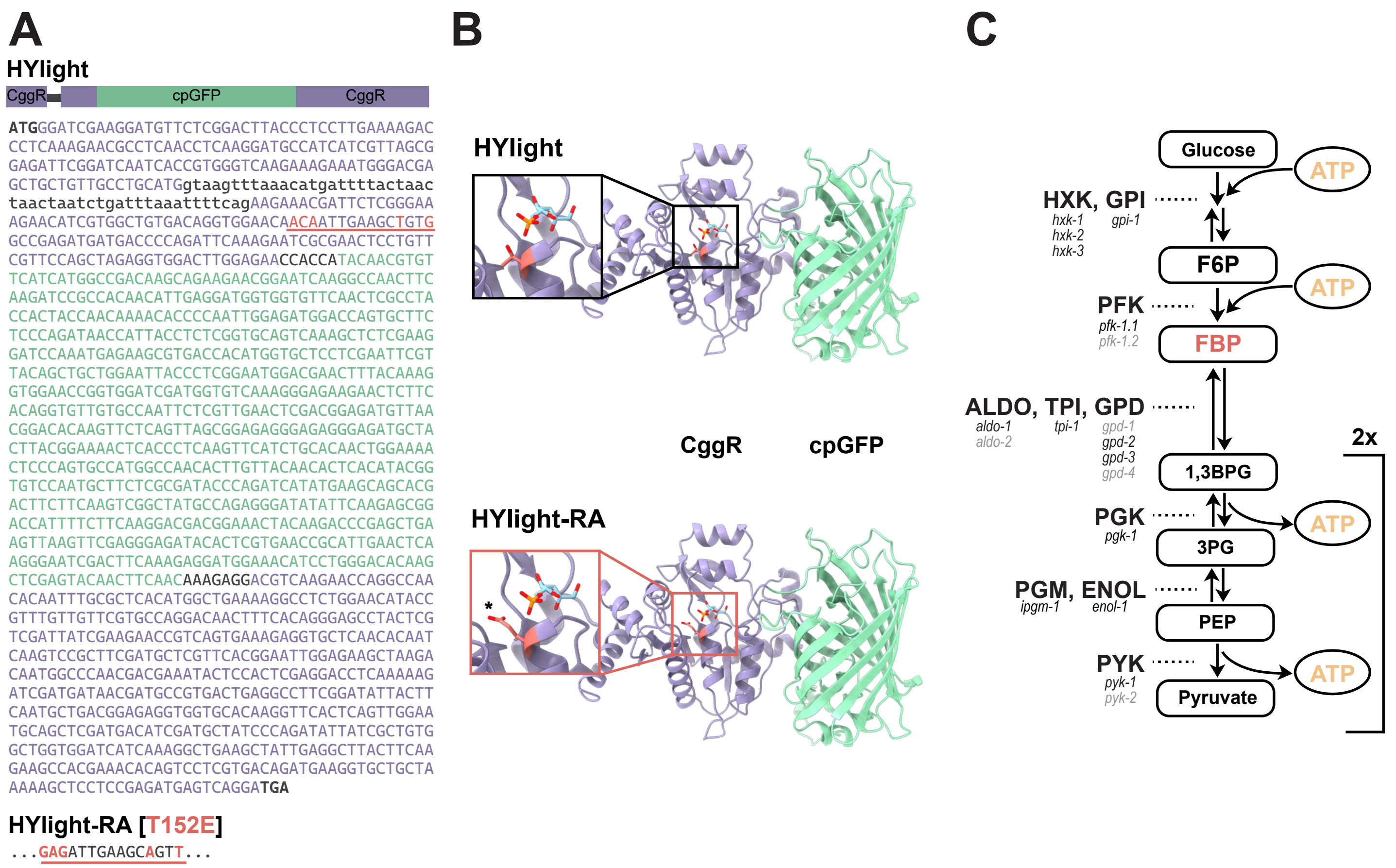


**Figure S1.** The HYlight biosensor measures FBP inside cells of *C. elegans*. **(A)** At top: diagram depicting the components of the sensor HYlight(1) adapted for expression in *C. elegans*. HYLight is comprised of a fusion between cpGFP and the Central Glycolytic Gene Regulator (CggR), a transcriptional regulator of metabolism from *Bacillus subtilis*. The CggR regions are shown in purple; cpGFP is shown in green. DNA sequence is shown below. The inserted intron at the 5' end is shown in lowercase bases. To generate HYlight-RA (bottom sequence), threonine 152 (in CggR numbering) is mutated to glutamate as shown in the red underlined region. This region also contains two silent mutations, which were added to improve cloning efficiency. **(B)** Structure models of HYlight compared to HYlight-RA. The model of HYlight was generated with AlphaFold2(2, 3), and the bound F6P molecule (see inset) was placed by aligning the crystal structure of CggR bound to FBP from RCSB ID 3BXF(4). The residue important for binding to FBP that is mutated in HYlight-RA is shown in red, with the glutamate mutation shown below (red inset). **(C)** Diagram of glycolysis, showing known homologs of each gene that are present in *C. elegans*. Genes that are less ubiquitously expressed in neurons based on scRNA-SEQ data(5) are shown in light grey. The location of FBP within the pathway is highlighted in red.

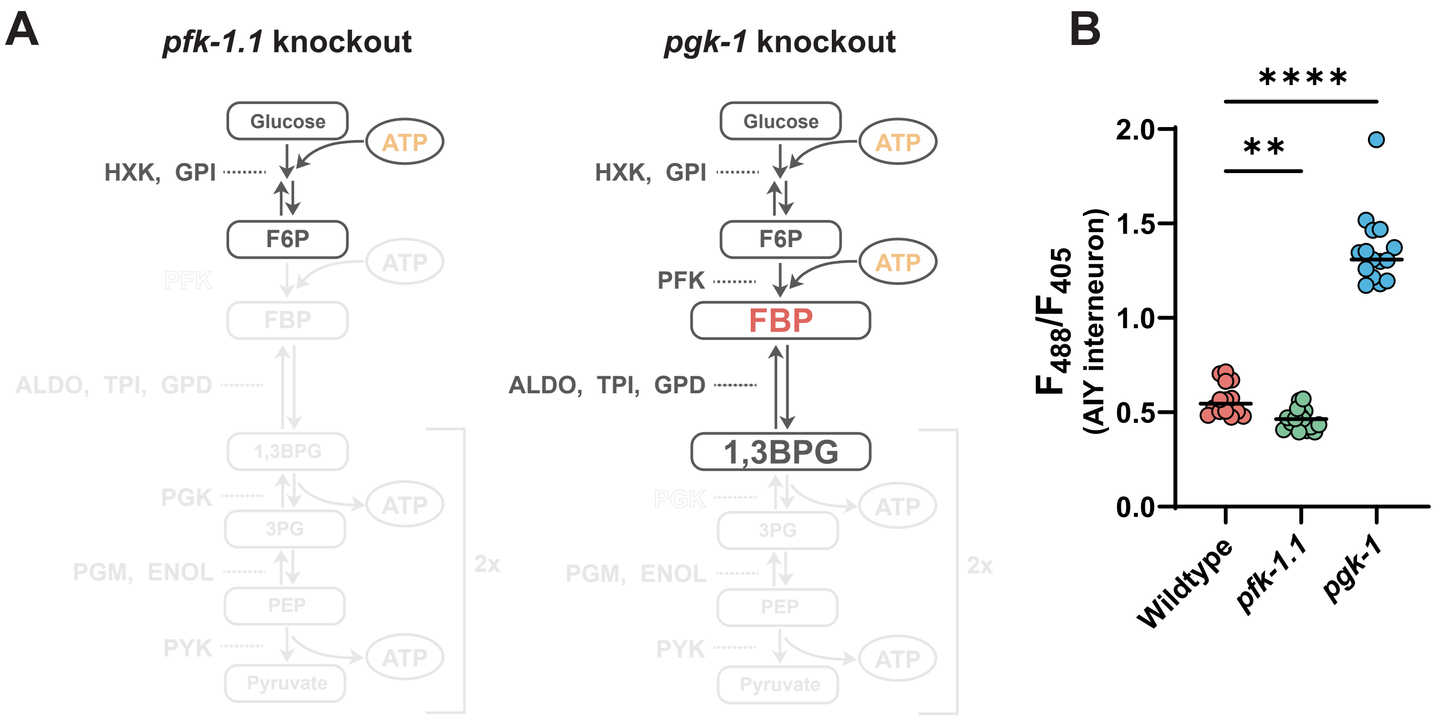


**Figure S2.** Genetic manipulation of FBP levels in *C. elegans*. **(A)** Schematic of the glycolytic pathway and expected affected steps for examined mutants. Mutation of the gene *pfk-1.1*, which catalyzes the conversion of F6P to FBP, should result in a near-complete elimination of FBP in cells (left), as they lack any alternative production pathway for the molecule. Conversely, loss of the gene *pgk-1*, which catalyzes a downstream reaction step converting 1,3BPG into 3PG, should result in an accumulation of FBP, as reduced usage would cause the reversible reactions between PFK and PGK to reach an elevated equilibrium value. **(B)** Measurement of HYlight ratios in the interneuron AIY for the indicated genetic backgrounds. Compared to wild-type worms, the *pfk-1.1(ola458)* mutant background results in a significantly reduced HYlight ratio (P-value: 0.0032) whereas the *pgk-1(tm1563)* mutant results in a significantly elevated ratio compared to wild type (P-value: <0.0001), as expected based on their position in the glycolytic pathway (A). Significance calculated by ANOVA with Tukey post-hoc test for multiple comparisons for 15 animals.

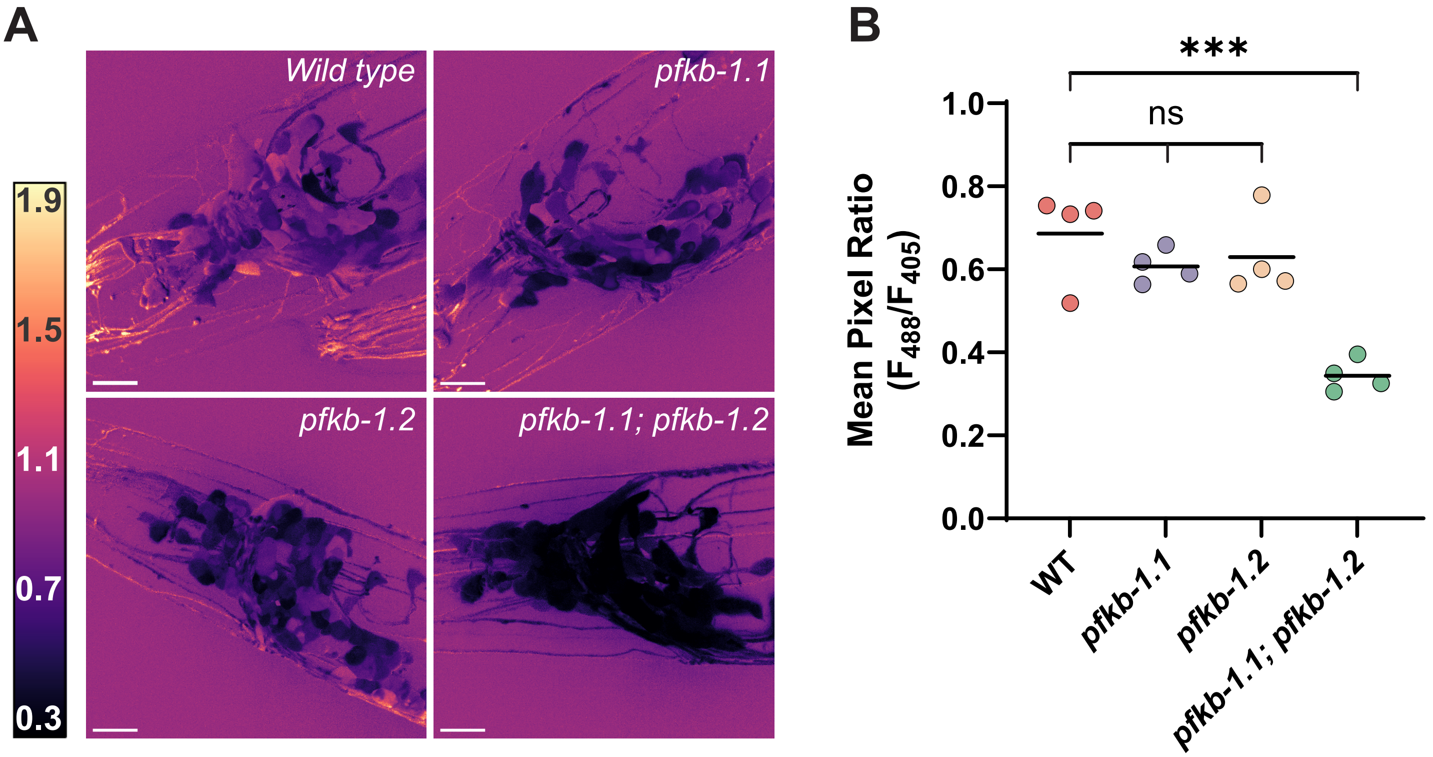


**Figure S3.** The two homologs of PFKFB, *pfkb-1.1* and *pfkb-1.2*, are functionally redundant in regulating neuronal glycolysis in *C. elegans*. **(A)** Images of pan-neuronal HYlight expression in either wild-type, *pfkb-1.1(ok2733)* mutant, *pfkb-1.2(ola508)* mutant or *pfkb-1.1; pfkb-1.2* double mutant backgrounds. Images of wild type and the double mutant are also shown in Figure 3 of this paper. Scale bars reflect 10 μm. **(B)** Quantification of HYlight values for indicated genetic backgrounds. Each dot represents individual animals. P-values shown are 0.529 (WT vs. *pfkb-1.1*), 0.750 (WT vs. *pfkb-1.2*), 0.980 (*pfkb-1.1* vs. *pfkb-1.2)*, and 0.0003 (WT vs. *pfkb-1.1; pfkb-1.2)*. Significance calculated by ANOVA with Tukey post-test for multiple comparisons across 4 individual worms of at least 60 Z slices.

**Table S1.** List of strains used within this study.

| Strain | Genotype | Source |
|---|---|---|
| N2 | Bristol wild-type strain | CGC |
| DCR3373 | *olaEx1951 [ttx-3p::tom-20:GFP; ttx-3p::mCherry:rab-3]* | (6) |
| DCR8881 | *olaEx5329 [rab-3p::HYlight]* | This study |
| DCR8892 | *olaEx5331 [rab-3p::HYlight-RA]* | This study |
| DCR8981 | *olaEx5367 [ttx-3p::HYlight]* | This study |
| DCR9016 | *pfk-1.1(ola458); olaEx5329* | This study |
| DCR9089 | *olaIs138 [ttx-3p::HYlight] IV* | This study |
| DCR9164 | *olaEx5443 [unc-47p::HYlight]* | This study |
| DCR9168 | *pgk-1(tm5613) I; olaIs138 IV* | This study |
| DCR9169 | *pfkb-1.1(ok2733) I; pfkb-1.2(ola508) IV; olaEx5367* | This study |
| DCR9197 | *olaIs138 IV; pfk-1.1(ola458) X* | This study |
| DCR9218 | *olaIs138 IV; pfk-1.1(ola458) X; olaEx5468 [ttx-3p::SL2::pfk-1.1]* | This study |
| DCR9238 | *pfkb-1.2(ola508) olaIs138 IV* | This study |
| DCR9251 | *pfkb-1.1(ok2733) I; olaIs138 IV* | This study |
| DCR9358 | *pfkb-1.1(ok2733) I; pfkb-1.2(ola508) IV; olaEx5329* | This study |
| DCR9359 | *pfkb-1.1(ok2733) I; olaEx5329* | This study |
| DCR9360 | *pfkb-1.2(ola508) IV; olaEx5329* | This study |
| DCR9421 | *pfkb-1.1(ok2733) I; pfkb-1.2(ola508) IV; olaEx5443* | This study |
| DCR9422 | *ric-7(n2657) V; olaEx1951* | This study |
| DCR9442 | *isp-1(qm150) IV; olaEx5367* | This study |
| DCR9443 | *olaIs138 IV; ric-7(n2657) V.* | This study |
| EG8040 | *oxTi302 I; oxTi75 II; oxTi411 unc-119(ed3) III; him-8(e1489) IV* | CGC |
| XE1763 | *oxTi76 IV; oxTi405 him-5(e1490) V; oxIs12 X* | Gift (M.H.) |

**Movie S1 (separate file).** HYlight response in neurons upon treatment with hypoxia. A single worm expressing pan-neuronal HYlight (*rab-3p)* was mounted on a microfluidics device (for applying transient hypoxia) as described in Methods. Frames were captured as 2-channel 25 µm Z stacks with 0.5 µm spacing every 30 seconds at 60x magnification. The first minute was captured in the presence of air; at time zero, the flow was switch to N2 gas, which then subsequently was captured for an additional 9 minutes.

**Movie S2 (separate file).** HYlight responses in AIY neurons upon treatment with hypoxia. Multiple worms expressing HYlight in the AIY neurons (*ttx-3p*) were imaged at lower magnification while experiencing transient hypoxia, as described in Methods. After an initial 1-minute treatment with air, a switch to N2 gas was performed (time 0) and images were captured for the remainder of the experiment in hypoxia. Images were captured every 5 seconds at a single Z plane at 10x magnification. Shown is a crop to 300 x 300 pixels with grayscale values for 405 excitation fluorescence on left and ratiometric HYlight (488/405) at right. Scale bar reflects 20 µm.
